# Proteasomal Dysfunction results in ER stress, Endo MT, oxidative stress, and apoptotic cell death resulting in Fuchs Corneal Endothelial Dystrophy like features in mice

**DOI:** 10.1101/2025.04.11.648416

**Authors:** Subashree Murugan, Sachin Anil Ghag, Viviane Souza de Campos, Hsuan-Yeh Pan, Marianne O. Price, Francis W. Price, Rajalekshmy Shyam

## Abstract

Fuchs Endothelial Corneal Dystrophy (FECD) is the irreversible degeneration of the corneal endothelium. The only treatment is corneal transplantation. To develop therapies for FECD, identifying the cellular causes for the onset and progression of the disease is crucial. While cell culture studies associate elevated oxidative stress, endoplasmic reticulum stress, endothelial-to-mesenchymal transition, and apoptosis with FECD, the causes behind the disease onset remain elusive. Guttae or Descemet’s membrane deposits are the earliest phenotype associated with FECD and are composed of unfolded proteins. Therefore, we asked if aberrant protein clearance pathways could be responsible for disease pathogenesis. We discovered a dysfunctional ubiquitin-proteasome pathway in a FECD mouse model and end-stage FECD patient samples. Inhibiting the ubiquitin-proteasome pathway in primary corneal endothelial cells resulted in the cellular dysfunctions associated with FECD. Finally, injecting healthy wild-type mice with proteasomal inhibitors resulted in all the major phenotypes associated with FECD, including corneal edema, guttae, and corneal endothelial cell loss. Therefore, this study strongly connects proteasomal dysfunction in FECD onset and progression.

## Introduction

The corneal endothelium, a post-mitotic cell monolayer on the posterior surface of the cornea, is responsible for maintaining the hydration and transparency of the cornea. Fuchs Endothelial Corneal Dystrophy (FECD) is a slowly progressive disorder affecting about 6 million Americans over 40 years of age (1–3). FECD is characterized by the presence of Descemet’s Membrane deposits (guttae), reduced corneal endothelial cell density, altered morphology, and corneal edema eventually causing poor visual acuity (4–6). Corneal transplant is the preferred treatment. While many genetic and environmental factors are associated with the disease onset (7–9), the initiation of the disease process is unknown.

Oxidative stress (7, 10), endoplasmic reticulum stress (11, 12), and endothelium-to-mesenchymal transition (13, 14) are all associated with FECD. The major sign of FECD, guttae, are made up of unfolded proteins (11, 15), which are a consequence of endoplasmic reticulum stress. Therefore, we asked if protein clearance pathways – autophagy and ubiquitin-proteasome pathway, responsible for removing accumulated proteins in the cells (16) - could be compromised during the disease onset. Whereas increased autophagy and mitophagy are associated with FECD (17), protein accumulation in the form of guttae persists. Therefore, we evaluated the status of the ubiquitin-proteasome pathway in FECD and its role in FECD onset and progression.

Using end-stage human FECD tissues, a newly developed mouse model for FECD (18), and *in-vitro* studies, we show that a dysfunctional ubiquitin-proteasome pathway is associated with the FECD signs. Finally, using wild-type animals, we show that pharmacological inhibition of proteasome activities is sufficient to induce FECD symptoms.

## Results

### Insufficient Ubiquitin Proteasome Pathway and upregulated autophagy in end-stage FECD samples

In FECD patient samples, we observed a significant decrease in the expression of Ubiquitin-Proteasome Pathway associated proteins - Proteasome 26S Subunit, Non-ATPase 11 protein (PSMD11), and the canonical signal for protein degradation by the proteasome K48-linked polyubiquitin and total poly-ubiquitin expression. These data indicate insufficient proteasomal activities in the end stage of FECD (**Fig. 1A and B**). We observed elevated expression of autophagosome marker, LC3II/I ratio, and decreased expression of autophagy substrate, Sequestosome (P62/SQSTM1) (**Fig. 1C and D),** revealing increased autophagy in FECD samples (17), consistent with previous findings.

**Figure 1:**
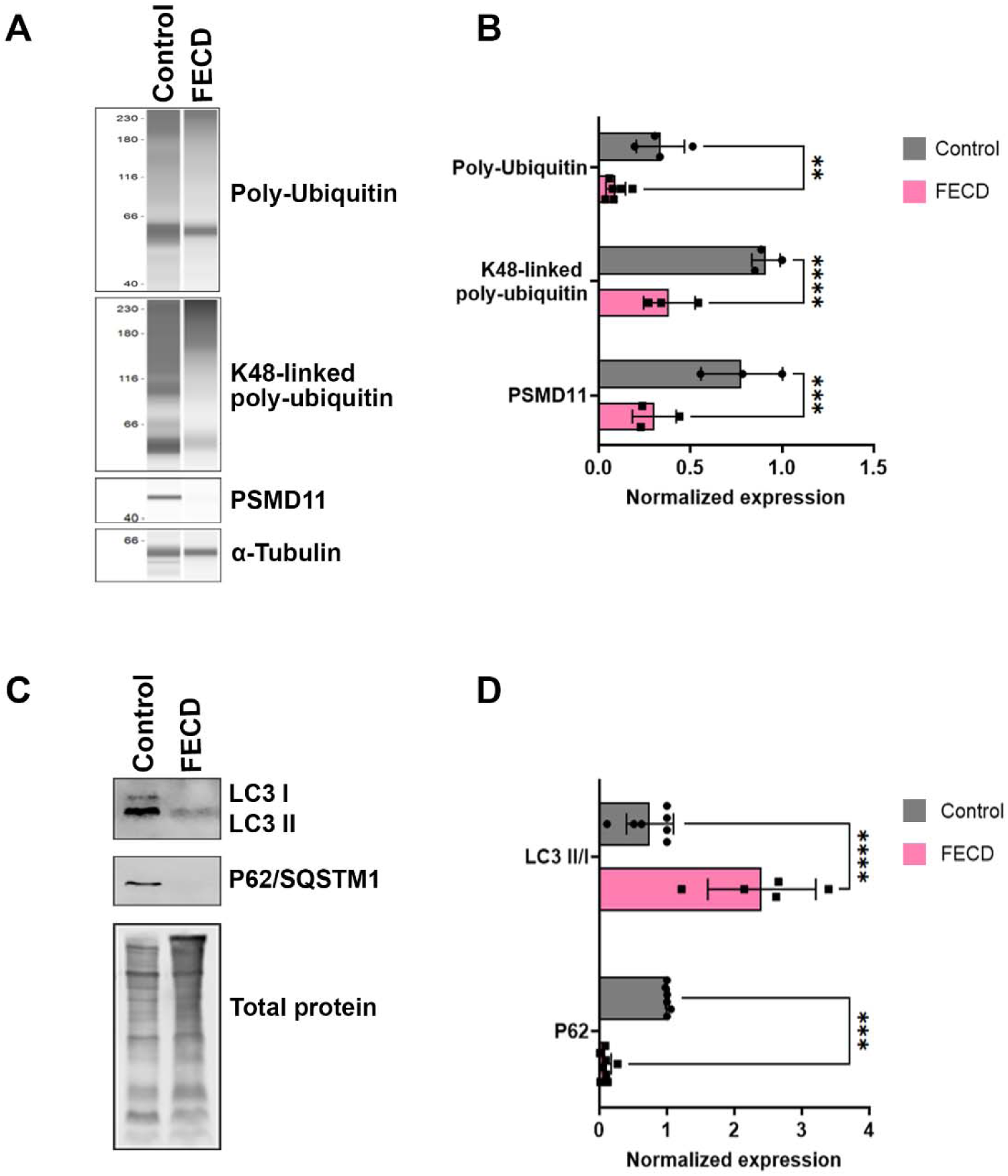
Decreased expression of Ubiquitin-Proteasome and increased expression of Autophagy pathway markers in end-stage human FECD samples. **A.** Representative blots from JESS immunoassay of FECD and age-matched control tissue samples for Poly-ubiquitin, K48-linked poly-ubiquitin, PSMD11 (Ubiquitin Proteasome Pathway markers), and α-tubulin. **B.** Quantification of the blots in panel A normalized to α-tubulin. N=3-6, Mean± standard deviation (SD) ** P<0.01, ***P<0.001, ****P<0.0001. (2-way ANOVA with Uncorrected Fisher’s LSD multiple comparisons). **C.** Representative traditional western blots of FECD and age-matched control tissue samples for LC3, P62, (Autophagy markers), and total protein. **D.** Quantification of the blots in panel C normalized to total protein. N=5-8, Mean± standard deviation (SD) ***P<0.001, ****P<0.0001. (2-way ANOVA with Uncorrected Fisher’s LSD multiple comparisons).

### Proteasomal inhibition results in FECD-associated dysfunctions in primary corneal endothelial cells

We tested whether proteasomal inhibition can induce FECD-associated cellular dysfunctions in corneal endothelial cells. Increased endothelial-to-mesenchymal transition (13), cell death (12, 19, 20), unfolded protein response (12), and oxidative stress (7, 10) are the major cellular features associated with FECD. To determine if proteasomal inactivity could induce these changes, we treated primary bovine corneal endothelial cell cultures with proteasomal inhibitor, MG-132. Primary cultures of bovine corneal endothelial cells are an established alternative to human corneal endothelial cells (21, 22). Unlike human corneal endothelial cells, these cells retain their endothelial characteristics and do not undergo endothelial-to-mesenchymal transition in cell culture (21, 22). As expected, we observed a 40% reduction in chymotrypsin activities in the MG-132 treated cells and an increase (∼15%) with betulinic acid, a proteasome activator, treatment (**Fig. 2A**).

**Figure 2:**
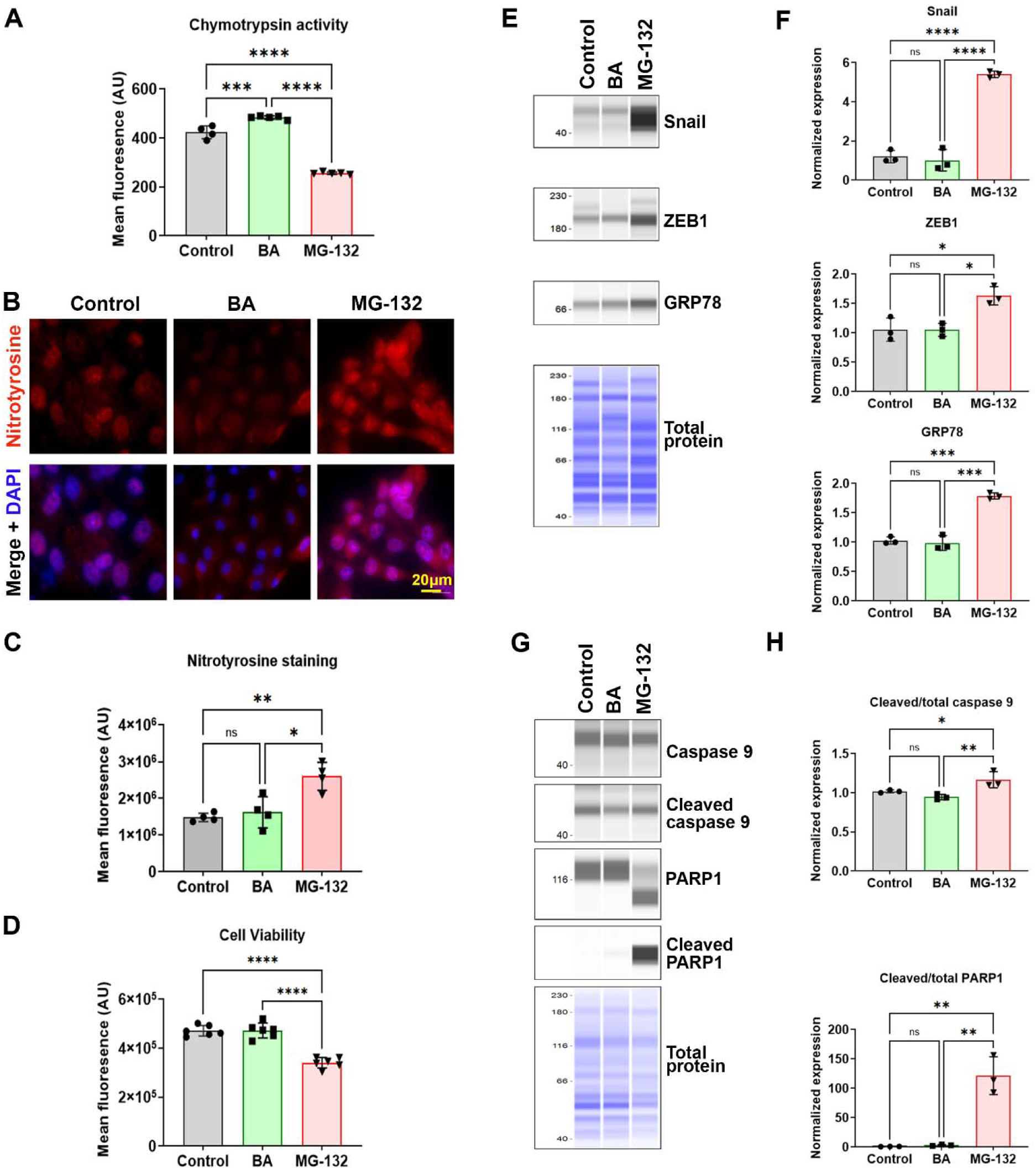
Proteasomal inhibition results in FECD-associated features in Bovine Corneal Endothelial Cells. **A.** Chymotrypsin activity of the 20S proteasome in cells treated with proteasomal activator (Betulinic acid, BA), proteasomal inhibitor (MG-132) and control. N=4-5, Mean± standard deviation (SD) ***P<0.001, ****P<0.0001. (2-way ANOVA with Tukey’s multiple comparisons). **B.** Nitrotyrosine staining of cells treated with BA, MG-132, and control. **C.** Quantification of the mean fluorescence from B. N=4, Mean± standard deviation (SD) *P<0.05, **P<0.01. (2-way ANOVA with Tukey’s multiple comparisons). **D.** Cell viability after treatments with BA, MG-132, and controls. N=6, Mean± standard deviation (SD) ****P<0.0001. (2-way ANOVA with Tukey’s multiple comparisons). **E.** Representative JESS immunoassay blots for endothelial-to-mesenchymal markers (Snail and ZEB1), Endoplasmic Reticulum stress marker (GRP78) and total protein in cells treated with BA, MG-132, and controls. **F.** Quantification of the blots in panel E normalized to total protein. N=3, Mean± standard deviation (SD) * P<0.05, ***P<0.001, ****P<0.0001. (2-way ANOVA with Tukey’s multiple comparisons). **G.** Representative JESS immunoassay blots for apoptosis markers (PARP1, cleaved-PARP1, caspase 9, cleaved-caspase 9) in cells treated with BA, MG-132, and controls. **H.** Quantification of the blots in panel G normalized to the total protein. N=3, Mean± standard deviation (SD) *P<0.05, **P<0.01. (2-way ANOVA with Tukey’s multiple comparisons).

MG-132 treatment showed a significant increase in nitrotyrosine, an oxidative stress marker (**Fig. 2B and C**), endothelial-to-mesenchymal transition markers, Snail, and Zinc finger homeodomain transcription factor (Zeb1). In addition, we also observed an increase in the expression of the master regulator of Unfolded Protein Response, Glucose-Regulated Protein, 78 kDa (GRP78) (**Fig. 2E and F**). A 27% decrease in cell viability (**Fig. 2D**) with an increase in the ratio of cleaved to total caspase 9 and PARP1 (apoptotic markers) was noted with MG-132 treatment (**Fig. 2G and H**). These in vitro experiments demonstrate that proteasomal inhibition can recapitulate key cellular dysfunctions observed in FECD, including increased oxidative stress, endothelial-to-mesenchymal transition, and apoptosis, highlighting the central role of proteasomal activity in maintaining corneal endothelial cell homeostasis.

### Proteasomal dysfunction is independent of unfolded protein response or oxidative stress in primary corneal endothelial cells

Two major factors associated with FECD pathogenesis are increased oxidative stress (10) and increased unfolded protein response (12)Therefore, using pharmacological approaches, we determined if proteasome dysfunction could be secondary to these distresses in primary corneal endothelial cells.

Thapsigargin is a potent inhibitor of sarco/endoplasmic reticulum Ca^2+^-ATPases, resulting in endoplasmic reticulum stress and unfolded protein response (23). Transcript levels of spliced X-box binding protein 1 (XBP-1) increase during unfolded protein response (23). Treatment with thapsigargin increased GRP78 protein levels and spliced XBP-1 mRNA expression (**Fig. 3A and B**). However, this treatment did not affect chymotrypsin protease activities (**Fig. 3C**) or proteasome-associated markers’ expressions (**Fig. 3D and E**). Consistent with previous findings, we found increased oxidative stress with thapsigargin treatment (**Fig. 3F and G**). The significant decrease in cell viability on thapsigargin treatment (**Fig. 3H**) corroborated the increase in the cleaved to total caspase 9 and PARP1 (**Fig. 3I and J**).

**Figure 3:**
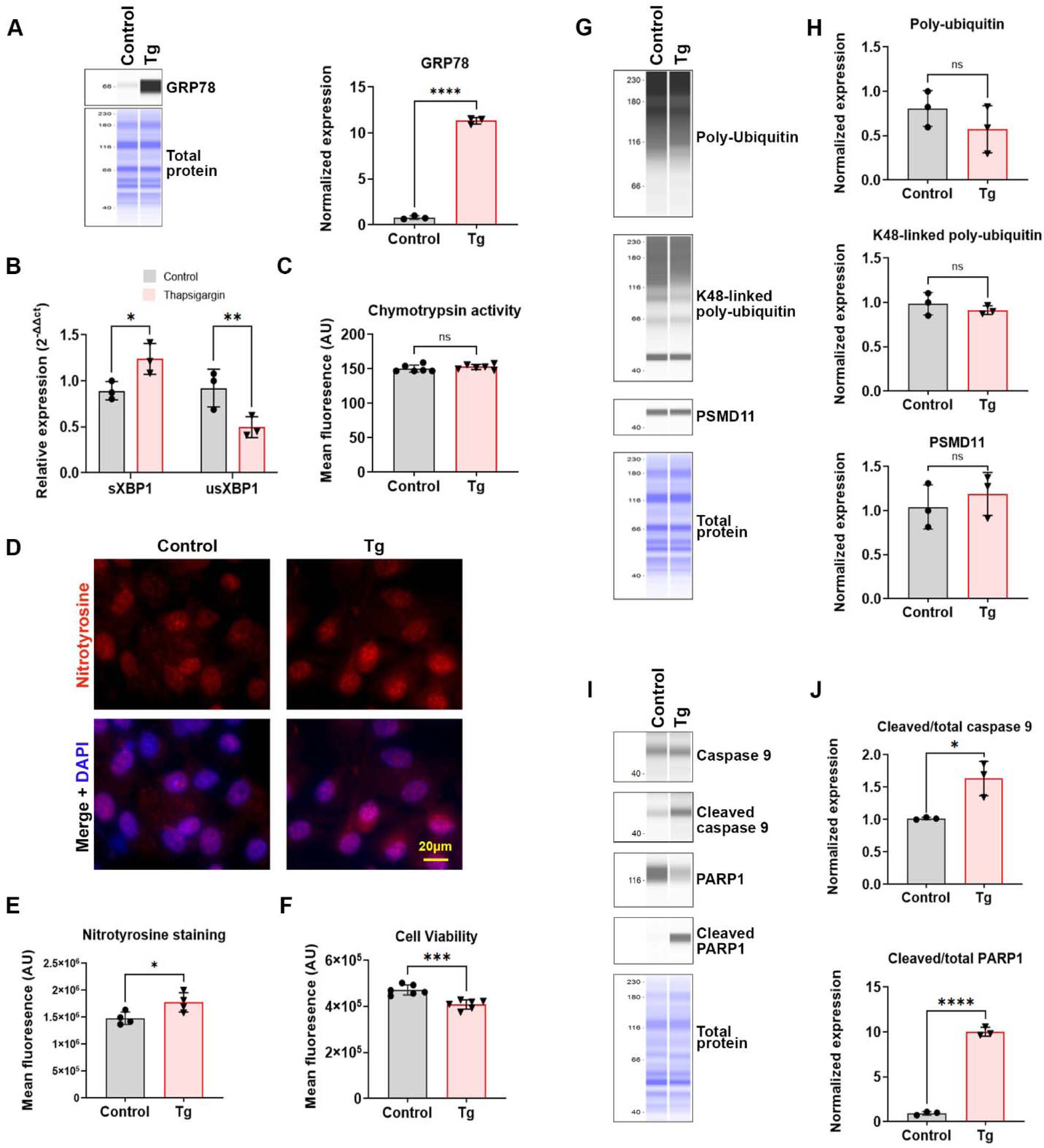
Effect of Endoplasmic Reticulum stress on FECD-associated features in Bovine Corneal Endothelial Cells. **A.** JESS immunoassay of GRP78 (Endoplasmic Reticulum stress marker) in thapsigargin treated and control cells and its quantification normalized to the total protein**. \*\***\*\*P<0.0001. (Unpaired t-test with Welch’s correction). **B.** Quantification of the expression of spliced and unspliced XBP1 transcripts in cells treated with Endoplasmic Reticulum stress inducer, Thapsigargin (Tg) and untreated controls. N=3, Mean± standard deviation (SD) * P<0.05, **P<0.01 (2-way ANOVA with Uncorrected Fisher’s LSD multiple comparisons) **C.** Chymotrypsin activity levels in thapsigargin treated and control cells. ns-not significant. (Unpaired t-test with Welch’s correction). **D.** Representative JESS immunoassay blots for Ubiquitin Proteasome Pathway markers (Poly-ubiquitin, K48-linked poly-ubiquitin, and PSMD11) and total protein in thapsigargin treated and control BCEC. **E.** Quantification of the blots in panel D normalized to the total protein. N=3, Mean± standard deviation (SD), ns-not significant. (Unpaired t-test with Welch’s correction). **F.** Representative nitrotyrosine staining with DAPI to visualize the reactive oxygen species in thapsigargin-treated and control cells. **G.** Quantification of the mean fluorescence from H. N=4, Mean± standard deviation (SD) *P<0.05. (Unpaired t-test with Welch’s correction). **H.** Quantification of the mean fluorescence indicative of the cell viability in cells treated with thapsigargin and controls. N=6, Mean± standard deviation (SD) ***P<0.001. (Unpaired t-test with Welch’s correction). **I.** Representative JESS immunoassay blots for apoptosis markers (PARP1, cleaved-PARP1, caspase 9, cleaved-caspase 9) in cells treated with thapsigargin and controls. **J.** Quantification of the blots in panel H normalized to the total protein. N=3, Mean± standard deviation (SD) *P<0.05, ****P<0.0001. (Unpaired t-test with Welch’s correction).

Tert-Butyl hydroperoxide (TBHP) induces oxidative stress (24, 25). We assessed whether acute or chronic oxidative stress can result in proteasomal dysfunctions. Treatment with 50 µM TBHP (24) (for 24 or 96 hours increased the mean fluorescence intensity of Nitrotyrosine staining (**Fig. S4A and B**). However, we did not observe changes in chymotrypsin activity (**Fig. 4C**). Although PSMD11 levels were slightly decreased after acute treatment there was no significant change in the ubiquitin-proteasome pathway-associated proteins on chronic exposure (**Fig. 4D and E**). Moreover, oxidative stress failed to increase endothelial-to-mesenchymal transition or Unfolded protein response after chronic exposure to TBHP (**Fig. 4F and G**). Consistent with previous reports (24, 25) there was increased apoptosis depicted by the increase in the cleaved-caspase 9 and cleaved-PARP1 expression (**Fig. 4H and I**) and decreased cell viability with TBHP treatment (**Fig. 4J**).

**Figure 4:**
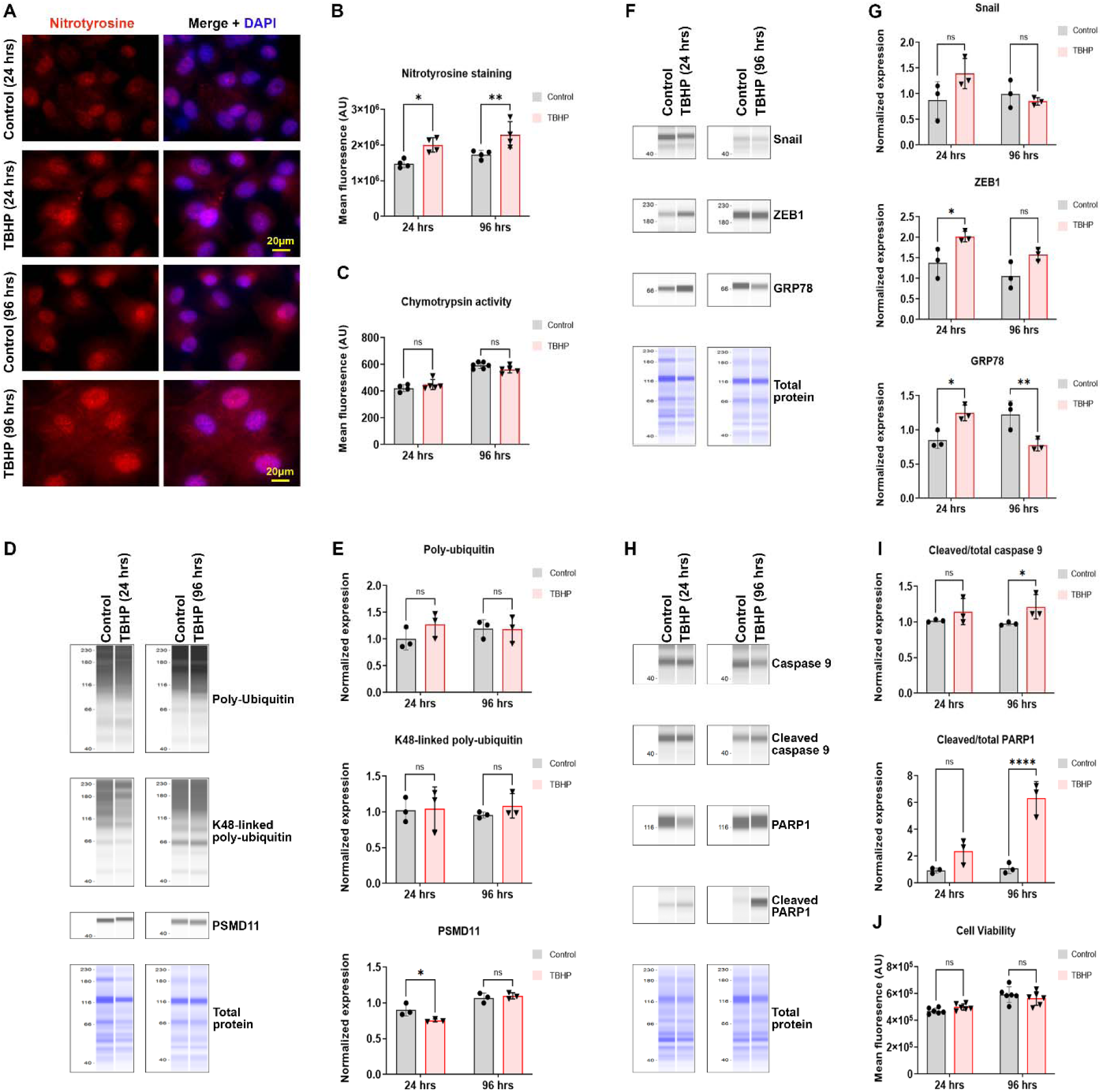
Effect of acute and chronic oxidative stress on FECD-associated features in Bovine Corneal Endothelial Cells. **A.** Representative nitrotyrosine staining with DAPI to visualize the reactive oxygen species in tert-butyl hydroperoxide (TBHP) treated (24 and 96 hours) and control cells. **B.** Quantification of the mean fluorescence from A. N=4, Mean± standard deviation (SD) *P<0.05, **P<0.01. (2-way ANOVA with Bonferroni’s multiple comparisons). **C.** Chymotrypsin activity of the 20S proteasome in TBHP treated and control cells. ns-not significant. (2-way ANOVA with Bonferroni’s multiple comparisons). **D.** Representative JESS immunoassay blots for Ubiquitin Proteasome Pathway markers (Poly-ubiquitin, K48-linked poly-ubiquitin, and PSMD11) and total protein in TBHP-treated and control cells. **E.** Quantification of the blots in panel D normalized to the total protein. N=3, Mean± standard deviation (SD), ns-not significant. *P<0.05 (2-way ANOVA with Bonferroni’s multiple comparisons). **F.** Representative JESS immunoassay blots for endothelial-to-mesenchymal markers (Snail and ZEB1), endoplasmic reticulum stress marker (GRP78) and total protein in TBHP treated and control cells. **G.** Quantification of the blots in panel F normalized to the total protein. N=3, Mean± standard deviation (SD) *P<0.05, **P<0.01, ns-not significant. (2-way ANOVA with Bonferroni’s multiple comparisons). **H.** Representative JESS immunoassay blots for apoptosis markers (PARP1, cleaved-PARP1, caspase 9, cleaved-caspase 9) in cells treated with TBHP (24 and 96 hours) and controls. **I.** Quantification of the blots in panel H normalized to the total protein. N=3, Mean± standard deviation (SD) *P<0.05, ****P<0.0001. (Unpaired t-test with Welch’s correction). **J.** Quantification of the mean fluorescence indicative of the cell viability in cells treated with TBHP and controls. N=8, Mean± standard deviation (SD) ****P<0.0001, ns-not significant. (2-way ANOVA with Bonferroni’s multiple comparisons).

### Proteasome inactivity coincides with the onset of disease phenotypes in a mouse model of FECD

Next, we assessed whether proteasome inactivity coincided with the onset of disease phenotypes in a newly developed FECD mouse model (18). We observed decreased polyubiquitin levels, K48 ubiquitylation, and increased PSMD11 expression at the onset of phenotypes in the FECD mice (**Fig. 5A and B**). We found a significant reduction in chymotrypsin activity in the corneal endothelia of the FECD mice (**Fig. 5C**).

**Figure 5:**
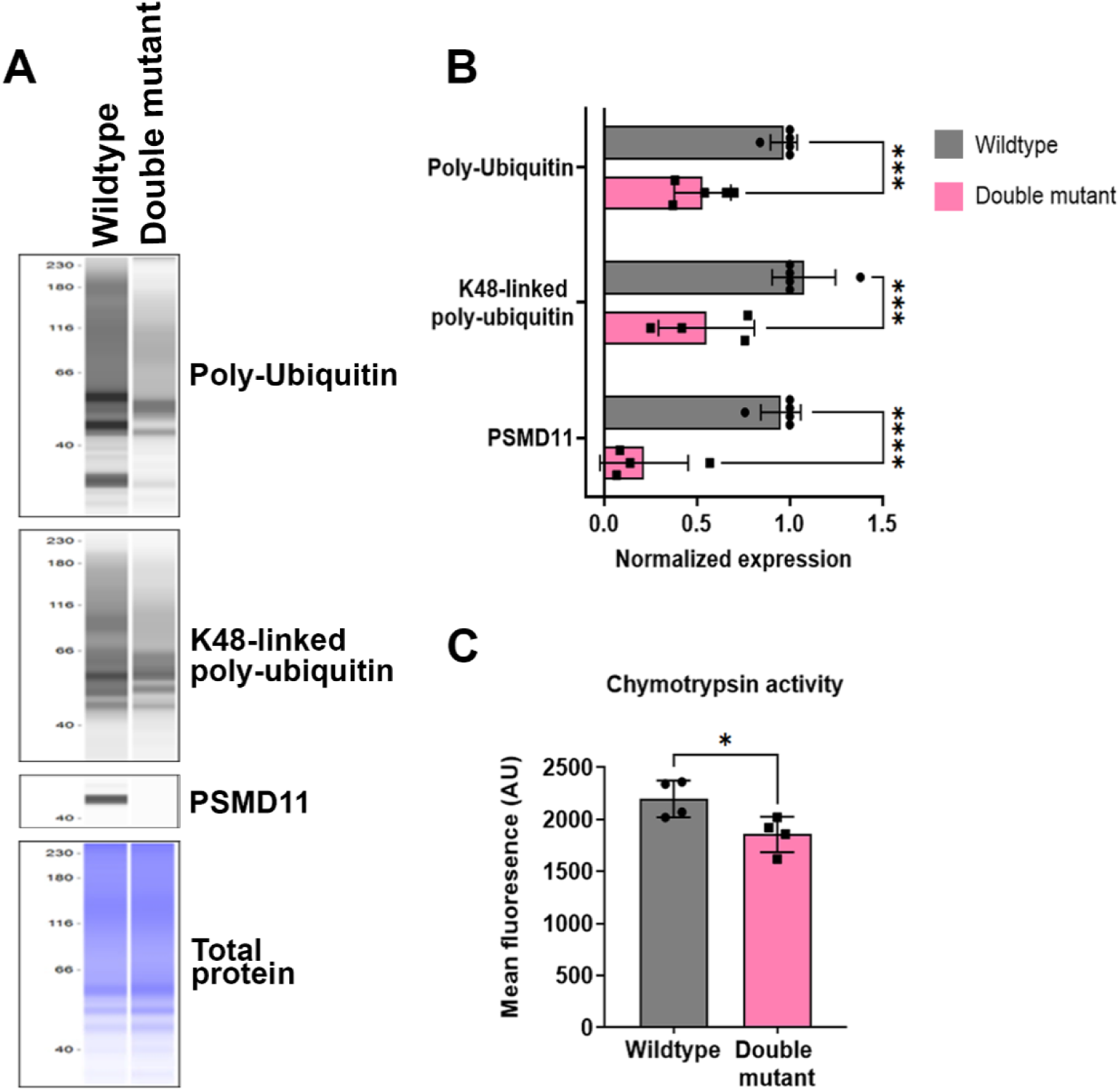
Ubiquitin-Proteasome pathway markers are downregulated in an FECD mouse model (*Slc4a11* knockdown and *Col8a2^Q455K^* knockin, or double mutant). **A.** Representative blots from JESS immunoassay of double mutants and age-matched wildtype animals for Poly-ubiquitin, K48-linked poly-ubiquitin, PSMD11 (Ubiquitin Proteasome Pathway markers) and total protein. **B.** Quantification of the blots in panel A normalized to total protein. N=3 to 5, Mean± standard deviation (SD) *** P<0.001, ****P<0.0001. (2-way ANOVA with Uncorrected Fisher’s LSD multiple comparisons). **C.** Chymotrypsin activity of the 20S proteasome in the corneal endothelium of wildtype and double mutant animals. N=4, Mean± standard deviation (SD) * P<0.05. (Unpaired t-test with Welch’s correction).

### Proteasome inactivity in wild-type mice induces FECD-like phenotypes

To test whether pharmacological inhibition of proteasome activities can result in FECD phenotypes, we injected 3-week-old C57BL/6J animals with proteasome inhibitors – MG-132 or Bortezomib. The major phenotypes associated with FECD include corneal edema, sub-endothelial guttae, endothelial cell loss, and changes in cell morphology. We evaluated the development of these features in the animals injected with proteasome inhibitors. The animals injected with Bortezomib, MG-132, and saline showed no significant difference in their body weights or any evident systemic defects (**Fig. 6A and B**).

**Figure 6:**
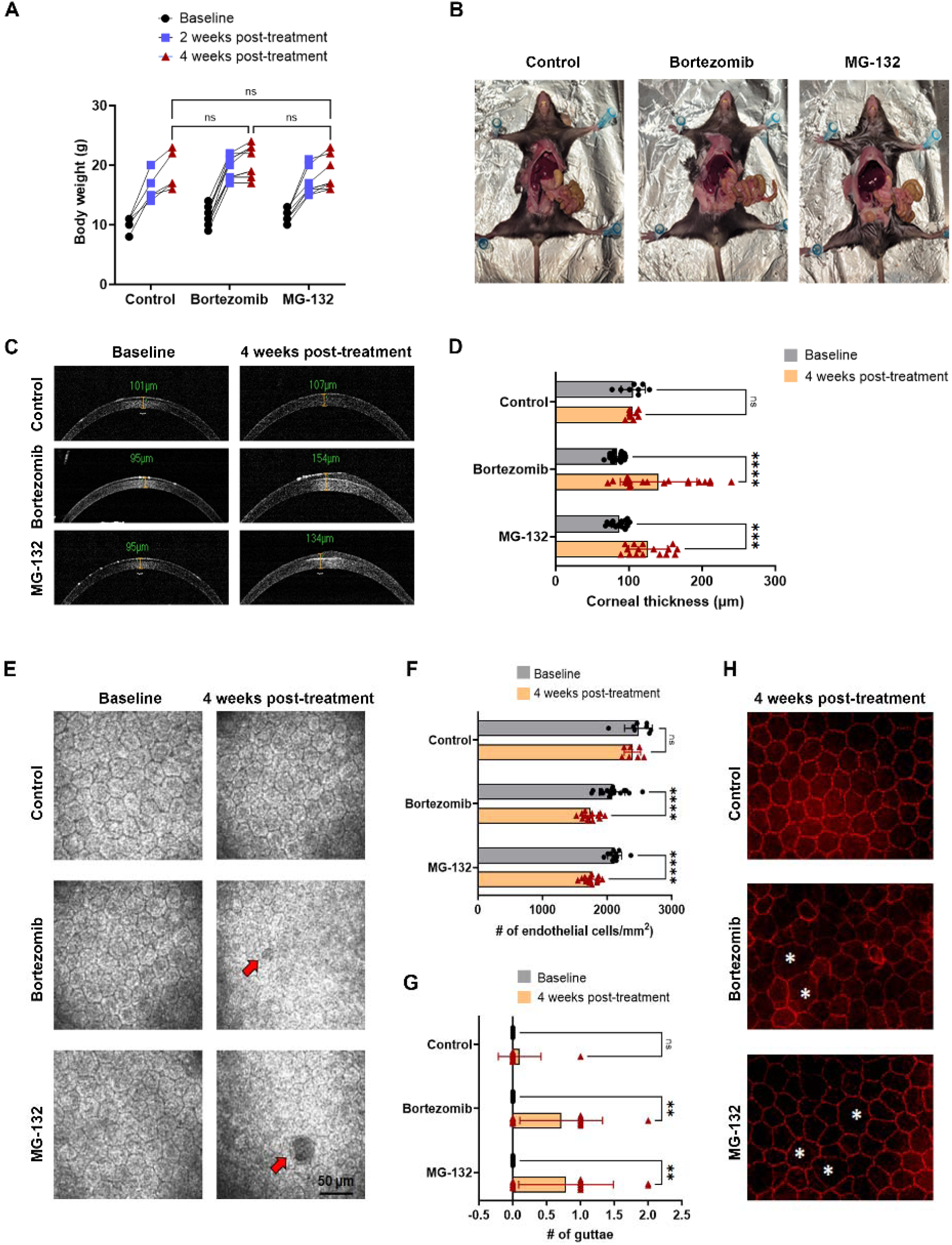
Altered corneal endothelium characteristics in wild-type mice after pharmacological inhibition of proteasomal activities. **A.** Longitudinal measure of the body weight in control, Bortezomib, and MG-132 injected animals at baseline, 2- and 4-weeks post-treatment. N=4 - 6, Mean± standard deviation (SD) ns-not significant. (2-way ANOVA with Tukey’s multiple comparisons). **B.** Representative dissections of control, Bortezomib, and MG-132 injected animals 4 weeks post-treatment showed no visceral abnormality. **C.** Representative OCT images of control, Bortezomib, and MG-132 injected animals at baseline and 4 weeks post-treatment. **D.** Quantification of the corneal thickness in the right and left eyes at baseline and 4 weeks post-treatment (n=8 to 18 eyes) **E.** Representative HRT3-RCM images of control, Bortezomib, and MG-132 injected animals at baseline and 4 weeks post-treatment (FOV 400µm), red arrows represent guttae. **F.** Quantification of the endothelial cell density/mm^2^ in the right and left eyes at baseline and 4 weeks post-treatment. N=8 to 18 eyes, Mean ± standard deviation (SD) ns-not significant, ****P<0.0001. **G.** Quantification of the number of guttae in the right and left eyes. N=10 to 14), Mean ± standard deviation (SD) ns-not significant, **P<0.01, ns-not significant. **H.** Representative images of ZO-1 staining of control, Bortezomib, and MG-132 injected animal whole corneas at 4 weeks post-treatment. Asterisks indicate polymorphism and polymegathism of the corneal endothelial cells.

Optical coherence tomography measurement revealed a significant increase in the corneal thickness (**Fig. 6C and D**), a substantial reduction in corneal endothelial cell density, and the presence of guttae in the MG-132 as well as Bortezomib injected animals (**Fig. 6E-G**). Consistently, we observed alterations in corneal endothelial cell shape and organization as evidenced by ZO-1 staining (**Fig. 6H**). We did not observe any significant changes in intraocular pressure (data not shown). All these data strongly suggest the involvement of proteasomal dysfunctions in developing the FECD phenotypes.

## Discussion

Here, we determined the role of ubiquitin proteasomal pathway dysfunctions in the development of FECD. Using a combination of *in-vitro* and *in-vivo* studies, we discovered that inhibiting the ubiquitin-proteasomal pathway is sufficient to induce many of the known FECD-associated dysfunctions (**Fig. 7**).

**Figure 7:**
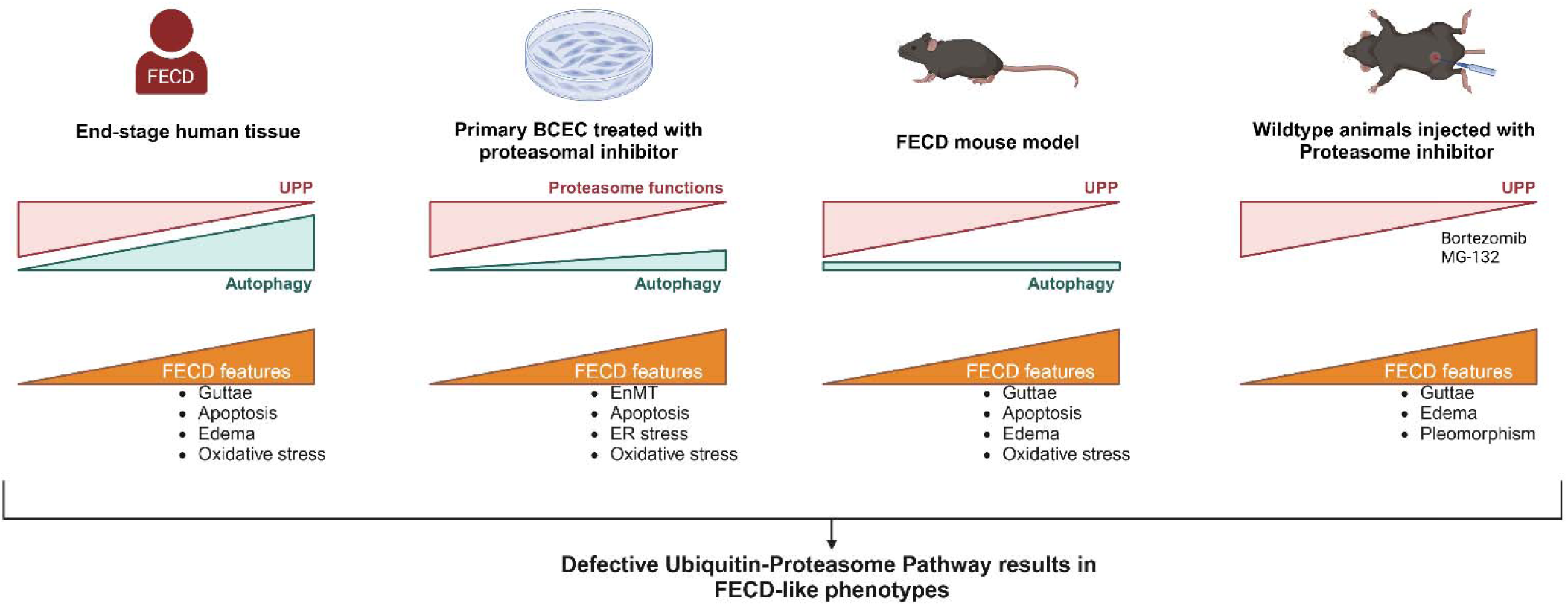
Graphical abstract of the findings. In this study we discovered defective ubiquitin proteasome pathway in end-stage FECD patient samples. Treating primary bovine corneal endothelial cells with proteasome inhibitors was sufficient to induce FECD associated cellular dysfunctions. In addition, we observed dysfunctional proteasome pathway at the onset of FECD features in a mouse model. Finally, inhibiting proteasome functions resulted in the FECD associated phenotypes in healthy, young wild-type animals.

Previous *in-vitro* studies show a strong association between oxidative stress (10, 15), endoplasmic reticulum stress, and endothelial-to-mesenchymal transition (12) with corneal endothelial dysfunctions. In addition, TCF4 intron expansion (26) and the female sex (27) are strongly correlated with FECD development. However, the causes leading to the disease onset remain unknown. In the current study, we provide evidence that inhibiting the ubiquitin-proteasome pathway can recapitulate the known cellular dysfunctions associated with FECD, such as elevated oxidative stress, endoplasmic reticulum stress, endothelial-to-mesenchymal transition, and cell death. Furthermore, pharmacological inhibition of the ubiquitin-proteasome pathway in young, healthy mice resulted in the major FECD phenotypes: corneal edema, corneal endothelial cell loss, and cell shape changes. These results strongly suggest that ubiquitin-proteasome pathway dysfunctions are sufficient to induce FECD phenotypes, opening the possibility of new therapies to slow disease progression.

While protein clearance pathways are crucial for cell health, their role in corneal endothelial cell functions is less understood. Previous studies uncovered dysfunctional autophagy, secondary to mitochondrial oxidative stress, in the *Slc4a11* KO mouse, a model for congenital hereditary endothelial dystrophy (28). Jurkunas *et al.* showed that end-stage FECD samples present elevated autophagy and mitochondrial dysfunctions (29). Dysfunctional autophagy was not determined to be causal to either FECD or Congenital Hereditary Endothelial Dystrophy. Whether elevated autophagy is a corneal endothelial cell adaptation to circumvent the insufficient proteasome activities in FECD is elusive.

*In-vitro* studies by Okumura *et al.* implicated ER stress associated unfolded protein response, and subsequent apoptosis in the corneal endothelial cells (11, 12). However, the effects of ER stress on oxidative stress, and guttae formation are not understood. In our study, ER stress induced elevated oxidative stress, and apoptosis, but failed to elicit proteasome dysfunctions. Similarly, oxidative stress associated mitochondrial dysfunctions and apoptotic cell death in corneal endothelial cell lines are well-established(15, 19, 25). However, oxidative stress failed to increase ER stress or proteasome dysfunctions in the present study. With these findings, we concluded that while ER stress and oxidative stress can result in apoptotic cell death, they alone cannot explain all the major cellular dysfunctions associated with FECD. However, proteasome inactivity increased oxidative stress, ER stress, endothelial mesenchymal transition and apoptotic cell death in primary corneal endothelial cells.

We developed the first mouse model for FECD that has all the disease-associated phenotypes(18). In this mouse model, we found decreased proteasome activities coincident with the onset of FECD phenotypes. However, at the age of analysis, we previously found elevated oxidative stress and decreased corneal endothelial cell density in these animals(18). These confounding factors limited our ability to determine the sufficiency of proteasome inactivity to induce FECD phenotypes in this mouse model. For this reason, we decided to pharmacologically inhibit proteasome functions in normal young mice right after weaning. FECD is an aging disease. We rationalized that if we can induce FECD like phenotypes in young, healthy mice, such an approach would eliminate the role of age, and other confounding variables in our analysis. Both bortezomib and MG-132 treatments resulted in corneal edema, guttae, and corneal endothelial cell loss; and recapitulated the three major phenotypes associated with FECD. These findings led us to conclude proteasomal inhibition to be sufficient to induce Fuchs Dystrophy like symptoms in-vivo.

Bortezomib is a chemotherapy agent used in the treatment of multiple myeloma. Ocular complications are rarely observed in patients undergoing this treatment. Chalazion and keratitis are common complaints among these rare instances(30, 31). While patients are often treated once a week with this drug for 18-30 days, our mouse experiments involved frequent administration at a higher dosage, which could explain the lack of corneal endothelial dysfunctions in the patients.

While our study provides evidence to support the role of proteasome inactivity in FECD pathogenesis, we do not have mechanistic insights on the causes leading to this. Similarly, we do not know how proteasome inactivity leads to oxidative stress, endothelial to mesenchymal transition, ER stress and cell death. Future studies to address these are on-going in our lab.

FECD is a blinding disease with no cure. By the year 2050, this disease is expected to affect 415 million people globally (32, 33). The only treatment currently available for this condition is corneal transplantation. With the aging population, the demand outweighs the supply of healthy donor tissue (15). Around the world, only one in 70 patients have access to the tissue (32, 33). Therefore, devising therapies to treat FECD is crucial. By determining the effects of the inactive ubiquitin-proteasome pathway in inducing FECD-like symptoms, this study provides novel insights into the disease pathogenesis.

## Materials and Methods

### Sex as a biological variable

Our study examined male and female human FECD tissue samples and animals. We report similar findings in both sexes.

### Human tissue

De-identified human corneal endothelial tissue removed from FECD patients during endothelial keratoplasty (a generous gift from Price Vision Group, Indianapolis) and age-matched control endothelium (purchased from Vision First, Indianapolis) were used for protein extraction as described previously (18).

### Transgenic mice

Mice containing tamoxifen-inducible knockdown of *Slc4a11* with the *Col8a2 (Q455K)* knock-in mutation were bred and maintained on tamoxifen chow as described previously (18). 16-week-old animals and age-matched wild-type animals were used for the experiments. Animal genotypes were carried out using automated genotyping services (Transnetyx, Cordova, TN).

### Bovine Corneal Endothelial Cell primary culture, pharmacological treatments, and capillary-based immunoassay

Corneal Endothelial cells were isolated from bovine eyes obtained from the local abattoirs. The cells were cultured in Dulbecco’s Modified Eagle Medium (DMEM) (Gibco, catalog # 11995065) containing 10% Fetal Bovine serum (FBS) (Gibco, catalog # 10082-147) and 1x antibiotic-antimycotic (Gibco, catalog # 15240062) until passage 2 and then used for experiments.(34) Cells were treated with Betulinic acid(35) (10 µM, Enzo Life Sciences, ALX-350-298), MG-132(36) (25 µM, Enzo Life Sciences, catalog # BML-PI102-005), thapsigargin(23) (0.25 µM, Millipore Sigma, catalog # 586006), tert-Butyl hydrogen peroxide(24) (50 µM, Sigma Aldrich, catalog # 458139) for 24 or 96 hours after overnight serum starvation (37, 38). The cells were washed twice with 1x Phosphate Buffered Solution (PBS) and lysed using 1x RIPA (Millipore Sigma, catalog # 20-188), 1x protease and phosphatase inhibitor (Cell Signaling Technology, catalog # 5872s), 25 µM MG-132 and 50 µM PR-619 (LifeSensors, catalog # SI9619). The lysate was centrifuged for 15 minutes (mins) at 13000x g at 4C. The supernatant was used for capillary-electrophoresis immunoassay as described previously (18, 28). The antibodies used were: Mouse anti-ubiquitin (Cell Signaling Technology, catalog # 3936s), Rabbit anti-PSMD11 (Cell Signaling Technology, catalog # 14303s), Rabbit anti-K48-linkage specific polyubiquitin (Cell Signaling Technology, catalog # 4289s), Rabbit anti-BiP (Cell Signaling Technology, catalog # 3177s), Rabbit anti-Snail (Cell Signaling Technology, catalog # 3879s), Rabbit anti-ZEB1 (Cell Signaling Technology, catalog # 70512s), Rabbit anti-Caspase 9 (Cell Signaling Technology, catalog # 9504s), Rabbit anti-cleaved-caspase 9 (Cell Signaling Technology, catalog # 9509s). Rabbit anti-PARP1 (Abcam, catalog # ab1912179504s), and Rabbit anti-cleaved-PARP1 (Abcam, catalog # ab32064).

### Traditional Western blotting

Protein lysates were denatured and contained 4x loading buffer with β-mercaptoethanol (Bio-rad, catalog # 1610710). 18 to 30 ug of protein were loaded and resolved in 12% or 15% SDS-PAGE. The bands were transferred to 0.22um nitrocellulose membrane (Li-COR, catalog # 926-31092) for 1 hour and blocked with blocking buffer (Li-COR, catalog # 927-60001) before incubating with primary antibodies – Rabbit anti-LC3 (R&D, catalog # MAB85582) and Mouse anti-P62 (Cell Signaling Technology, catalog # 88588ss) diluted in antibody diluent (Li-COR, catalog #65001), overnight at 4C. Incubation with rabbit or mouse secondary antibody (Li-COR, catalog # 926-68071 or 926-32210) was performed for 1 hour at room temperature. The blots were imaged using Odyssey XF imager (Li-COR, catalog #2802-01). Total protein stain (Li-COR, catalog #926-10016), to measure the total protein in each sample, was used for normalization.

### Nitrotyrosine staining

The passage 2 bovine corneal endothelial cells were grown up to 80% confluence in an 8-chamber slide and treated with Betulinic acid, MG-132, thapsigargin, and tert-Butyl hydrogen peroxide for 24 or 96 hours after overnight serum starvation. Nitrotyrosine staining was performed as previously described (18). The cells were washed with 1x PBS twice and fixed using 100% methanol for 10 mins. They were permeabilized and blocked using 0.5% Triton X-100 (Fisher Scientific, catalog # BP151-500) and 5% Normal donkey serum (Jackson Immuno Research, catalog #017-000-121) in 1x PBS for 30 mins to 1 hour at room temperature followed incubation with Rabbit anti-Nitrotyrosine (Life Technologies Corporation, catalog #A21285) (1:100) overnight at 4°C. The cells were washed with 1x PBS thrice and then incubated with Goat anti-Rabbit IgG; Alexa Fluor™ 488 (Thermo Fisher, catalog #A11034) secondary antibodies at 1:100 dilution, at room temperature for an hour. The cells were mounted using Prolong Glass Antifade Mountant containing NucBlue (Invitrogen, catalog #P36985). The cells were imaged with a Zeiss Apotome2 microscope (Carl Zeiss, White Plains, NY, USA). There were at least 4 replicates for each treatment and all images were obtained with consistent microscope settings for laser intensity, gain, and exposure time. ImageJ was used to analyze the captured images. The fluorescence intensity of at least 5 cells from each replicate was measured and the mean intensity per square area was plotted.

### Proteasome activity assay

The chymotrypsin-like protease activity was measured using a commercially available Amplite Fluorometric Proteasome 20S activity assay kit (AAT Bioquest, catalog # 13456) with LLVY-R110, a fluorogenic indicator that produces green fluorescence when cleaved by the proteasome. The mouse tissue or primary cells were incubated with the working solution for 1 hour at 37°C per the manufacturer’s instructions. For mouse tissue, corneal flat mounts were placed on a 96-well plate with the endothelium facing up. The fluorescence was measured using a microplate reader at 520-530 nm with 480-500 nm excitation.

### Cell viability assay

The number of viable bovine corneal endothelial cells after treatment with pharmacological agents was evaluated using CellTiter-Glo 2.0 Assay (Promega, catalog # G9241). This assay measures the ATP levels in the metabolically active cells reported as luminescence. The manufacturer’s instructions were carried out as follows: After 24 or 96 hours of treatment in a 96-well plate, the plate was equilibrated to room temperature for 30 mins. An equal volume (100 µL) of working reagent was added to the wells, contents mixed for 2 mins using a shaker and incubated at room temperature for 10 mins to stabilize the luminescent signal. The signal was recorded using a microplate reader with an integration time of 0.3 seconds/well. A decrease in the luminescence is indicative of cell death.

### RNA extraction, reverse transcription, and real-time PCR

The bovine corneal endothelial cells treated with Thapsigargin were collected for RNA extraction using Trizol method. The cells were homogenized in 200 µL of Trizol reagent (Ambion, catalog # 15596018) using a motor and pestle. For phase separation, 40 µL of chloroform was added and mixed vigorously by hand for 15 seconds. The samples were incubated for 5 mins at room temperature and centrifuged at 13000x g for 15 mins at 4°C. The aqueous phase was transferred to a new tube and an equal volume of isopropanol was added and mixed by inversion 6-8 times. Further incubated at room temperature for 10 mins and centrifuged at 13000x g for 15 mins at 4°C. To pellet the RNA out, 1 mL of 75% ethanol was added and vortexed for 5 seconds, centrifuged at 13000x g for 15 mins at 4C, decanted the supernatant, and air-dried the pellet. The pellet was resuspended in 15 µL of the Nuclease-free water. The RNA was purified using the RNeasy Mini kit (Qiagen, catalog #74104) manufacturer’s protocol.

The cDNA was synthesized as previously described using High-Capacity RNA-to-cDNA kit (Applied Biosystems, catalog #4387406).(18) Briefly, the reaction mix contained 10 µL of the 2x RT buffer mix, 1µL of 20x RT enzyme, total RNA of 1µg and nuclease-free water up to 20 µL per reaction and incubated at 37°C for 1 hour. The reaction was halted by heating the samples at 95°C for 5 mins.

Real-time PCR was performed using SSO Advanced Universal SYBR green supermix (catalog #1725271) following the manufacturer’s protocol. The primers for bovine spliced XBP1 (sXBP1): Forward - GCTGAGTCCGCAGCAGGT and Reverse - CTGGGTCCAAGTTGAACAGAAT and unspliced (usXBP1) Forward - CAGACTACGTGCACCTCTGC Reverse - CTGGGTCCAAGTTGAACAGAAT were used.

### Injections, Optical Coherence Tomography, and Heidelberg Retinal Tomography 3

Wild-type animals (C57BL/6J, # 000664) purchased from Jackson lab were injected intraperitoneally with proteasome inhibitors Bortezomib (Hikma Pharmaceuticals, catalog # NDC 0143-9098-01) or MG-132 (Cayman chemicals, catalog # 13697) at 3-weeks of age. 0.5mg/kg of Bortezomib (39) dissolved in saline, was injected twice a week, and 15 µM/kg of MG-132(40) dissolved in DMSO, was injected 3 times a week for 4 weeks. Animals injected with saline thrice a week for 4 weeks were used as controls. Anterior segment-Optical Coherence Tomography (AS-OCT) (iVue100 Optovue, Inc., Fremont, CA, USA) and Heidelberg Retinal Tomography 3-Rostock Cornea Module (HRT3-RCM) (Heidelberg Engineering Inc., Franklin, MA, USA) were performed to measure the corneal thickness and assess the corneal endothelial cells, every two weeks from baseline up to 4 weeks post-treatment as previously described (18).

### ZO-1 staining

The corneal cups dissected from the mouse eyes were rinsed with 1x PBS and transferred to a 96-well plate for ZO-1 staining. The corneal cups were fixed using 4% paraformaldehyde in 1x PBS at room temperature for 10 mins, washed twice with 1x PBS, blocked and permeabilized using a buffer containing 0.5% Triton X-100 and 5% Normal donkey serum in 1X PBS for 30 mins to 1 hour at room temperature. The Mouse anti-ZO-1 (Fisher Scientific, catalog #339100) (1:100) antibody was used for primary incubation at 4°C overnight. The corneal cups were washed in 1x PBS and incubated with Goat anti-Mouse IgG, Alexa Fluor™ 594 (Thermo Fisher, catalog #A11032) for an hour at room temperature. Radial cuts were made in the corneal to mount them flat using Prolong Glass Antifade Mountant with NucBlue. The flat mounts were imaged using Zeiss Apotome2 microscope.

### Statistical analysis

All the data were quantified and analyzed using the GraphPad Prism program (v.10.3.1, Boston, MA, USA). The mean and standard deviation were represented using dot plots with error bars. Statistical significance and comparison were performed using one-way analysis of variance (ANOVA) for more than 2 groups or t-test for 2 groups (p-value less than or equal to 0.05 was considered statistically significant).

### Study approval

All human tissue samples were obtained from an ophthalmologist or a technician from the eye bank, after surgical removal. The researchers had no direct contact with the patient or the donor family. All animal experiments were performed following the Institutional Animal Care and Use Committee (IACUC) guidelines and the current regulations of the National Institute of Health (NIH) and the Association for Research in Vision and Ophthalmology (ARVO) statement for the use of animals in ophthalmic and vision science research.

## Data availability

All relevant information about raw data is available directly from the corresponding author.

## Acknowledgments

We are grateful for the support from Ms. Alejandra Rosas and Ms. Shreyasi Ganguly for their expertise with animal work and Dr. Darcy Trader Jones’s (UC Irvine) guidance throughout this study. We completed this project with funding support from the National Institute of Health R00 EY032974.

